# DORA: a dose-response autoencoder for interpretable transcriptome-to-viability prediction

**DOI:** 10.64898/2026.05.27.728125

**Authors:** Shuhui Wang, Alexandre Allauzen, Vaitea Opuu, Philippe Nghe

## Abstract

Predicting the effect of drugs on cell viability is a central challenge in drug discovery. Artificial intelligence holds the promise to considerably accelerate this process by leveraging rich cellular data such as transcriptomics. Current models focus on either transcriptomes or inhibitory concentrations, but they fall short in integrating these sources of information. Here, we propose DORA (Dose-Response Autoencoder), a deep learning model that predicts changes in transcriptomes and viability in a dose-dependent manner, knowing the unperturbed cell state. By enforcing a latent space consistent with cumulative dose effects, DORA matches other methods at predicting transcriptomes and substantially outperforms existing latent representations at viability prediction. The transcriptome-viability relationship provided by the model further allows the recovery of known biomarkers of cell viability while suggesting novel ones. Overall, DORA provides a unified framework delivering actionable biological insights for phenotypic drug screening and personalized medicine.

## 1 Introduction

Identifying potent and safe drugs requires extensive experimental campaigns to explore the vast chemical space of candidate compounds, with a low success rate: 1% from hit to lead, among which approximately 10% are finally approved after clinical trials [1, 2]. A major hope for drug discovery is that artificial intelligence could predict cellular responses to perturbations [3], thus allowing the early elimination of inefficient or toxic drug candidates [4]. Multimodal cell state modeling is a promising avenue for such a task. In this area, transcriptomics and phenotypes are expected to be mutually informative. Indeed, cells with similar transcriptomes are likely to respond similarly to drugs. Additionally, it might be possible to identify the subsets of genes that specifically respond to certain drugs, as well as those that cause phenotypic alterations. However, integrating these layers of information remains an open challenge.

The most scrutinized phenotype is cellular viability, as it is used to detect drugs that inhibit tumor cell growth and to quantify toxicity. Early computational approaches for viability prediction focused on chemical structure, namely Quantitative Structure-Activity Relationship (QSAR) methods. However, these initial approaches had limited success [5, 6], notably because they ignore cell type. Indeed, later studies improved IC50 predictions using machine learning strategies that jointly represent transcriptomes of untreated cells and drug descriptors [7, 8].

The rise of RNA-seq technologies has pushed our ability to characterize transcriptomic changes in bulk batches and single-cells (scRNA-seq) in response to drugs. LINCS-L1000 is one of the largest bulk transcriptomic perturbation datasets, profiling over 10^6^ signatures across more than 70 cell lines and over 1,000 drugs. Sci-Plex captures transcriptomics responses at the single-cell level, across 3 cell lines and 188 compounds. Together, these datasets provide complementary views of drug action: bulk RNA-seq offers breadth across drugs and conditions, while scRNA-seq reveals heterogeneous responses at the single-cell level.

Recent efforts focused on predicting transcriptomic responses with deep learning (DL). Latent space models (e.g., autoencoders) compress high-dimensional transcriptomics data into lower-dimensional representations while preserving perturbation signals [9, 10]. CPA [11] is an autoencoder that decomposes the perturbed transcriptomics signal into independent attributes for the cell and the drug. To enforce the independence of these attributes, CPA uses the method of adversarial training, which is, however, notoriously difficult to stabilize [12]. Besides, because each attribute embedding is learned purely from data, CPA requires substantial training data to yield reliable latent representations [13]. Building on CPA, Chem-CPA incorporated graph neural networks (GNNs) to improve drug molecular embeddings and thereby generalize to unseen compounds [14]. These methods are based on a latent space, where the representation of each attribute is combined additively. Biolord [15] proposed a different strategy. Instead of the additive latent space approach, the learned latent attributes (molecular and genetic) are concatenated and fed to the decoder, allowing them to combine nonlinearly. This improves training stability but inflates the latent dimensionality, reducing interpretability. Instead of disentangling features in the latent space, other approaches, such as CellOT and CellFlow model perturbations as transformations of trajectories of gene expression [16, 17], or distributions thereof in the case of single-cell transcriptomics.

To date, there are fewer attempts to combine transcriptomics and viability in the same model. Indeed, there is no sufficiently large dataset where both were measured at once across cells and drugs. Instead, multiple datasets must be aggregated. Viability measurements campaigns include CTRP, NCI60,or DrugComb [18, 19, 20]. For instance, DrugComb contains viability measurements for ∼8,000 drugs across *>*2,000 cell lines [20]. Some have built cross-referenced datasets, by identifying the overlap of drugs and cell lines between LINCS-L1000 transcriptomics and CTRP cell viability datasets [21, 22]. For instance, Szalai et al. [21] obtained an intersection of ∼4,000 (cell, drug) pairs, starting from two datasets of ∼ 6 × 10^6^ transcriptomics perturbations and 6 × 10^6^ viability measurements. They applied models such as Linear Regression and XGBoost, using transcriptomics as input and cell viability as output. They report improved viability prediction compared to using only drug descriptors, confirming the insight that transcriptomics informs on how cells respond to drugs.

Supposing that we have a model able to predict drug-dose-transcriptome-viability relationships, clinical decision-making then requires extracting restricted gene panels that can serve as biomarkers. In the case of gene expression, biomarkers are genes whose expression level provides criteria for diagnostics and guide of therapeutic interventions [23, 24]. Differential-expression (DE) analysis followed by pathway enrichment is the standard method to identify candidate biomarkers [25, 26]. DE analysis maps to each gene a single coefficient, facilitating the biomarker identification. Doing so, however, ignores an explicit dose structure and misses cooperative or antagonistic genetic interactions [27]. Closing the gap between computational methods and clinical practice requires models that (i) learn multigene interaction effects, (ii) encode dosage explicitly, and (iii) assign clear coefficients to individual genes, enabling functional validation.

Here, we introduce DORA (DOse-based Response Autoencoder), a latent space model designed to predict cellular viability and identify biomarkers using transcriptomics perturbation. The first distinctive feature of our approach is that DORA explicitly models the variation of drug doses as an iterative trajectory in its latent space. With this design, DORA achieves state-of-the-art performance in the prediction of gene perturbations on the LINCS and Sci-Plex datasets. We then use the latent representation obtained with DORA to achieve viability prediction, outperforming the available DL latent representations. The second distinctive feature of DORA is its constrained linear decoder architecture. It enables us to attribute clear viability coefficients to genes, facilitating biomarker identification. These genes were found to overlap with known cancer-inhibitor biomarkers. With its compact architecture and its accuracy, DORA offers a compelling alternative in-between large DL models and DE analyses.

## 2 Model and data

### 2.1 DORA model

Drug treatments induce complex, high-dimensional gene expression responses that vary with both dosage and cell type, but only a subset of features reflects meaningful changes in cellular state. A suitable method should extract these relevant, dose- and cell-type-dependent patterns by learning a low-dimensional representation. Recent datasets, such as LINCS, measure gene expression after different drug and dose treatments, providing a perfect foundation for developing such a method. To this end, we introduce the Dose-Response Autoencoder (DORA). DORA is designed to model transcriptomic changes relative to an untreated control under progressively increasing drug doses. As illustrated in Fig. 1a, the model learns a latent representations that encode the transition from the untreated control condition, denoted *d*_0_, to higher treatment levels, up to the maximum dose *d*_*n*_. By construction, the latent representation of the control condition is anchored at zero for each cell type, reflecting an absence of drug-induced perturbation.

**Figure 1:**
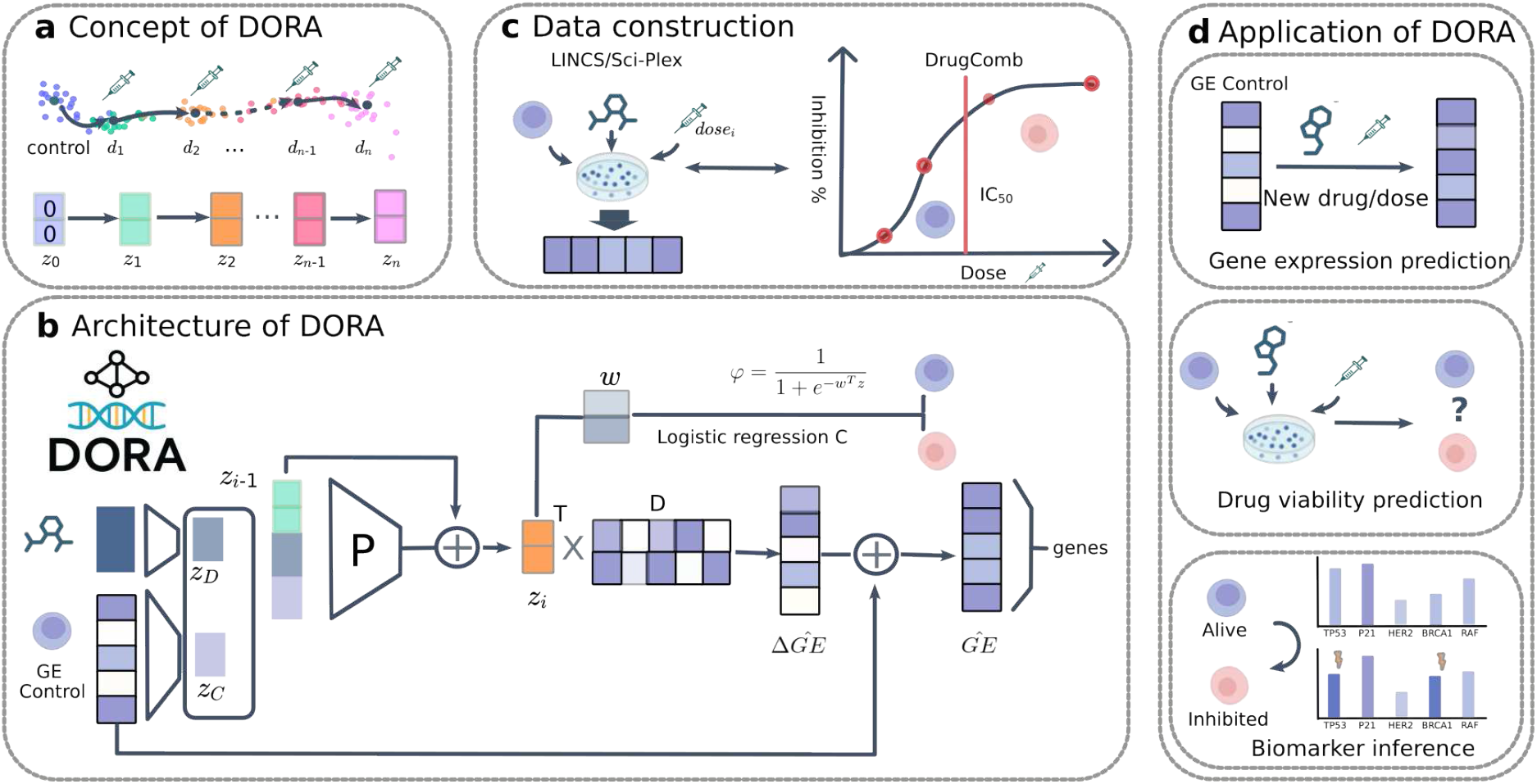
DORA for inference of transcriptomic data and cell viability prediction. **a**, Conceptual framework of DORA: We hypothesize that perturbations in the latent space are dependent on prior states, where distinct latent states correspond to unique cell phenotypes. **b**, Model architecture: DORA is a deep generative model for transcriptional response prediction comprising a forward latent module (P), a decoder (D), and a classifier (C). The model initializes with *d*_0_ (a vector of zeros); drug, dosage, and cell representations are input to module P, which models latent-space perturbations induced by drugs and cells. Sequential addition of multiple P modules models the trajectory of drug administration with increasing doses. Decoder D infers gene expression, while latent representations *z*_*i*_ are fed into classifier C to predict cell viability. **c**, Data reconstruction: To align gene expression and phenotypic data, we used gene expression data from the LINCS and Sci-Plex datasets, and drug response data from DrugComb; overlapping measurements for the same (cell, drug) combinations were extracted. **d**, DORA applications: (1) Gene expression prediction: Given control gene expression, DORA predicts gene expression following administration of a drug at a specified dose; (2) Drug viability prediction: Given control gene expression, drug, and dose, DORA predicts cell viability; (3) Biomarker identification: DORA is interpretable, enabling identification of the most highly perturbed genes associated with changes in cell viability.

DORA consists of three main modules (Fig. 1b): (i) The forward latent module learns a trajectory conditioned on the cell type and drug, using an iterative updating scheme of latent states, which reflects incremental changes in the cellular response to increasing drug dosages; (ii) the decoder module maps each point along this latent trajectory to a perturbed gene expression profile; and (iii) the phenotype module maps the latent representation to a phenotype, here cell viability.

Specifically, the input to DORA consists of individual tuples (C, D, *d*), where C is a cell line, D is a drug,and *d* is a dosage level. Each cell line is encoded by an embedding *z*_*C*_, computed from its non-perturbed gene expression profile *GE*(*d*_0_), and each drug is represented by an embedding *z*_*D*_ from a pretrained BERT model on drug SMILES (ChemBERT2a [10], see Section 7 for details).

The latent trajectory begins from a fixed initial state *z*_0_ = (0, 0, · · ·, 0) in *R*^*m*^. The forward latent module, implemented as a three-layer multilayer perceptron (MLP), iteratively computes latent states using residual updates *z*_*i*_ = *f* (*z*_*i*−1_, *z*_*C*_, *z*_*D*_) + *z*_*i*−1_, up to the step corresponding to dosage *d*_*n*_, producing *z*_*n*_. Although the model does not take the dosage value as an explicit input, increasing dosage is modeled implicitly: each application of the forward module corresponds to one step along the latent trajectory, so that each iteration corresponds to an implicit increase of concentration determined by a dose schedule unified across all drugs. To this aim, gene expression profiles for each cell line–drug pair are organized as dose-ordered sequences:

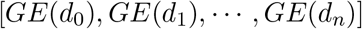

where *GE*(*d*_*i*_) denotes the gene expression profile measured at dosage level *d*_*i*_, and the doses satisfy *d*_0_ *< d*_1_ *<* · · · *< d*_*n*_.

The decoder module is a linear transformation parameterized by a matrix *D* ∈ *R*^*m*×*N*^, which maps *z*_*i*_ to the predicted gene expression profile 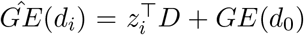. The phenotype module is a linear logistic regression model that maps *z*_*i*_ to the predicted viability 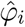.

We train two versions of the model. The first version, denoted DORA, is trained to reconstruct transcriptomic trajectories: at each dose step, the predicted expression profile is compared with the observed profile using a mean squared error loss. The second version, denoted DORA_*ft*_, uses the trained DORA and fine-tunes it jointly with the phenotype classifier, using only the samples with viability annotations (see Data construction section). The classifier receives the latent state and outputs the predicted viability probability. The total loss combines the reconstruction error with a binary cross-entropy loss for viability prediction. All model parameters are optimized jointly using the Adam optimizer [28]. For viability, we report both the predictions DORA and DORA_*ft*_.

### 2.2 Dataset construction

To train the model, we need data comprising both gene expression data and cell viability in response to increasing doses of drugs. However, no such data has been acquired in a single consistent dataset. Instead, we cross-referenced genomic data with cell inhibition information from distinct datasets, as done in Szalai et al. [21]. We combined RNA-seq data from LINCS (bulk) and Sci-Plex (scRNA-seq) with monotherapy drug response data from DrugComb [20]. LINCS comprises 28 cancer cell lines treated with 320 drugs, whereas Sci-Plex includes 3 cancer cell lines and 188 drugs. To assign viability labels (growth-inhibited vs. alive), we identified overlapping drug–cell line pairs across RNA-seq and DrugComb (see Fig. 1c). DrugComb provided dosage-response data for only 3–5 doses per drug, cell pair, so we first fitted a sigmoid model to each inhibition curve to reconstruct the level of inhibition for doses along a continuous scale (see Section 7). Samples were categorized as alive or inhibited based on their viability. After integration and filtering: i) the LINCS-derived dataset included 4,895 samples, with 12.2% inhibited; and ii) the Sci-Plex-derived dataset included 648 samples, with 14.4% inhibited.

This integrated dataset supports training and evaluation of (Fig. 1d): (1) predicting transcriptional responses at unobserved dosages or for unseen drugs, (2) estimating drug efficacy in terms of viability, and (3) identifying biomarkers associated with cellular response.

## 3 Prediction of drug-induced perturbations

To test whether the learned representation and model architecture capture the variation observed between cell types, drugs, and dosage, We evaluated DORA on the task of reconstructing gene expression. Experiments were conducted on both bulk RNA-seq data from the LINCS dataset and single-cell RNA-seq data from the Sci-Plex dataset, as seen in Fig. 2a. For evaluation, we split the data into 80% training, 10% validation, and 10% test sets. The split is performed at the trajectory level, ensuring that all dosage points from a given cell-drug pair are assigned to a single split. This prevents information leakage between training and evaluation. Performance is measured by computing the coefficient of determination (*R*^2^) between predicted and observed perturbed gene expression profiles. We considered two evaluation settings: (i) random split, where points on trajectories are assigned randomly to each subset; (ii) unseen compounds split, where entire drugs are held out during training to test generalization to unseen treatments; and (iii) unseen cell split, where cell lines are held out to evaluate generalization across cellular contexts.

**Figure 2:**
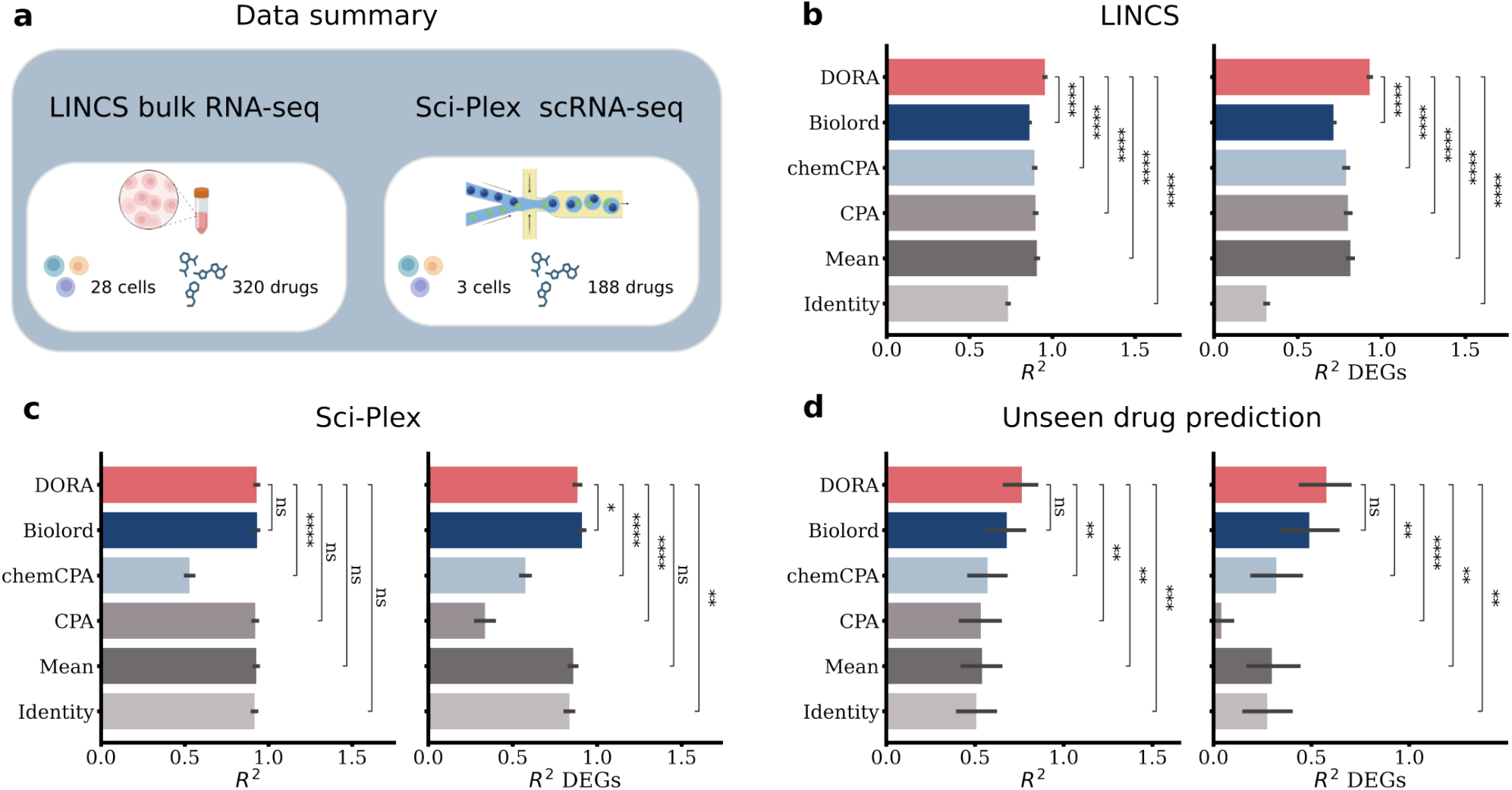
Prediction of gene expression (GE) using DORA and other state-of-the-art models (CPA, chem-CPA, Biolord) on the LINCS and Sci-Plex datasets. **a**, Data summary: LINCS consists of bulk RNA-seq data spanning 28 cell lines and 320 drugs; Sci-Plex comprises scRNA-seq data covering 3 cell lines and 188 drugs. **b**, Performance under the random-split scenario on LINCS. **c**, Performance under the randomsplit scenario on Sci-Plex. **d**, Performance under the unseen-compounds-split scenario on Sci-Plex. Left box plots show *R*^2^ scores across all genes; right box plots show scores for the top 50 DEGs. Results are presented as mean values ± SD. t-test is used to assess if the performances between models are statis-tically significant. ns: 0.05 *< p* ≤ 1, *: 0.01 *< p* ≤ 0.05, **: 0.001 *< p* ≤ 0.01, ***: 0.0001 *< p <*= 0.001,****: *p* ≤ 0.0001.

We compared DORA with five baselines. First, we considered three state-of-the-art autoencoder-based models: CPA, chemCPA, and Biolord. CPA models gene expression as a linear combination of one-hot encoded factors. chemCPA extends CPA by replacing the one-hot drug encoding with a graphbased representation. Biolord, in contrast, learns disentangled representations of biological factors and combines them nonlinearly to predict gene expression under perturbation. Second, we compared DORA with predictions consisting in averaging gene expression profiles computed from the training set. This baseline quantifies the extent to which models capture perturbation-specific effects beyond global mean expression patterns, denoted Mean. Finally, as a lower-bound reference, we included a null model that assumes no transcriptional response to treatment and returns the unperturbed (control) gene expression profile, denoted Identity. This baseline establishes a minimal performance threshold for assessing reconstruction accuracy. The detailed description for these models can be found in Section 7.

For random splitting in LINCS, DORA achieved an average *R*^2^ = 0.92, outperforming CPA (*R*^2^ = 0.86, with t-test *p <* 10^−4^), the Mean model (*R*^2^ = 0.86 with t-test *p <* 10^−4^), and all other baselines. The overall high performance of the models is in fact due to the moderate average effect of drugs, as indicated by the null model with *R*^2^ = 0.79. To better disentangle differences in model performances, we restricted the analysis to the top 50 differentially expressed genes (DEGs) (as defined in [11]). In this case, DORA still outperformed the other baselines (*R*^2^ = 0.93), see Fig. 2b. On the Sci-Plex dataset, DORA, Biolord, and CPA achieved comparable performances (*R*^2^ = 0.93, *R*^2^ = 0.93, *R*^2^ = 0.92 respectively), but the high score of the null model *R*^2^ = 0.92 suggests weaker drug perturbations overall. Unexpectedly, chemCPA underperformed, yielding *R*^2^ scores below the null model. We hypothesize that the adversarial training may require data-specific fine-tuning to converge. When restricted to the top 50 DEGs, DORA and Biolord significantly outperformed baselines (t-test with *p* = 0.02). In particular, CPA failed to accurately predict DEGs (see Fig. 2c). Furthermore, reconstruction accuracy on the Sci-Plex dataset (*R*^2^ ∼ 0.92) is observed to be higher than that on the LINCS dataset (*R*^2^ ∼ 0.86). As illustrated in Fig. SI1, this discrepancy arises from inherent differences between the datasets: the perturbation effect in Sci-Plex is notably weaker relative to that in LINCS.

We also evaluated performance on differential gene expression profiles, which isolate the drug-induced component of the response by removing gene expression of control (detail see Section 7); DORA again achieved the best results, with *R*^2^ reaching 0.66, significantly better than the other models (*p <* 10^−4^, see Fig. SI2). These results demonstrate that DORA captures well the important features of the dosedependent gene expression perturbations.

For the unseen compounds splitting, we selected 9 drugs not seen during training at the 4 dosage levels 0.01, 0.1, 1.0 and 10.0 *µM*, following previous work [11, 14, 15], namely: Dacinostat, Givinostat, Belinostat, Hesperadin, Quisinostat, Alvespimycin, Tanespimycin, TAK-901, and Flavopiridol. At low dosages, drug-induced perturbations are typically weak, while at high dosages, stronger responses are expected but are harder to predict accurately. At the highest dosage (10 *µM*), as seen in Fig. 2d, DORA achieved an average *R*^2^ = 0.80, comparable to Biolord (*R*^2^ = 0.79), outperforming CPA and chemCPA. For the top 50 DEGs, DORA consistently outperformed all state-of-the-art methods. For all models, predictions exhibited high variance across drugs, suggesting a high biological heterogeneity in drug perturbations. For instance, Flavopiridol induced strong transcriptional shifts for K562 and MCF7 cell lines but not for A549 (*R*^2^ = 0.92 for the highest dosage). As expected, model performance generally declined at higher dosages, likely due to strong responses that fall outside the range observed during training, see Fig. SI3a. While DORA uses a drug representation similar to the other state-of-the-art methods, it extrapolates more effectively across drugs.

For the unseen cell splitting, we held out one cell line as validation, and repeated this for all cell lines in the Sci-Plex data. Among the cell types tested (A549 for lung, K562 for leukemia, and MCF7 for breast), K562 proved to be the most challenging, with the lowest performance (*R*^2^ = 0.33 at 10 *µm*), compared to A549 (*R*^2^ = 0.69) and MCF7 (*R*^2^ = 0.52) (see Fig. SI3b). The differences in prediction quality are also consistent with hematopoietic cells having distinct transcriptional responses than the epithelial-derived A549 and MCF7 lines.

These findings indicate that DORA learns a latent representation able to capture key variation in drug responses, structured by dosage and cellular context. Below, we test whether the resulting embeddings support generalization to viability prediction.

## 4 Viability prediction from dose-structured latent space

We showed above that the representation learned by DORA captures the main sources of variation in gene expression data, reflecting drug- and dose-dependent effects. We hypothesized that this representation may be consistent with the mapping from genotype to phenotype. In this section, we test this by evaluating the prediction of cell viability from the latent representation computed on gene expression profiles.

To evaluate latent representations, we extracted latent representations from CPA, chemCPA, Biolord, and DORA, each pre-trained on the LINCS and Sci-Plex datasets (see Fig. 3a). For each latent representation, we trained a separate neural network classifier composed of three hidden layers with ReLU activations: the input dimension depends of the latent representation of the model considered (*e*.*g*. 32 for DORA on Sci-Plex and 128 on LINCS), while the second layer has half of its dimension, which is ultimately processed by a single neuron to output a probability using a sigmoid activation function. The classifier is trained on 70% of the labeled data (randomly selected) for 100 epochs using each representation. Viability prediction performance is evaluated with the remaining 30% of the labeled data using the area under the precision-recall curve (PR-AUC). We compared the latent space representations with the raw gene expression profiles and the PCA (top-100 components) representation of the gene expression.

**Figure 3:**
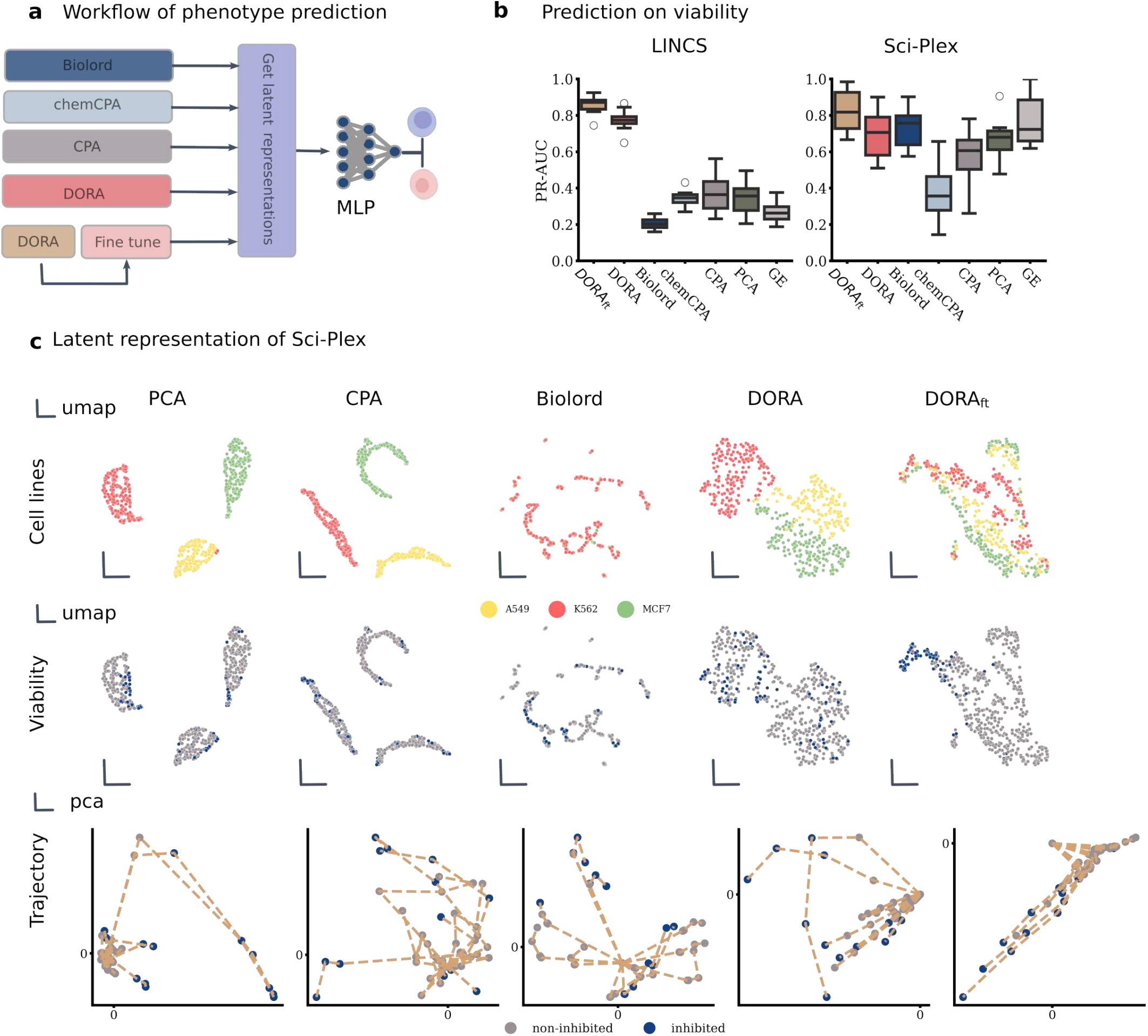
Comparison of models for phenotype prediction and latent representation analysis. **a**, Workflow: Latent representations from different models were used as input to a multi-layer perceptron (MLP) for cell viability prediction. **b**, Cell viability prediction performance (PR-AUC) on the LINCS and Sci-Plex datasets using original GE, PCA-reduced GE, or latent embeddings from CPA, chemCPA, Biolord, and DORA. DORA was additionally fine-tuned to assess performance gain (DORA_*ft*_). Box plots show the distribution of results from 10-fold cross-validation. **c**, Visualization of latent representations of Sci-Plex: Low-dimensional representations generated by PCA, CPA, Biolord, DORA, and fine-tuned DORA. UMAP representations are provided for cell lines and viability, with trajectories displayed for 8 randomly selected drug–cell pairs. Each trajectory starts from the control condition and extends to the highest dose; grey dots indicate non-inhibited samples, and blue dots indicate inhibited samples.

For the viability prediction task, the latent representation extracted from DORA on the LINCS dataset outperformed all other pre-trained models, achieving a PR-AUC of 0.77, significantly better (*p <* 0.05) compared with the second best representations obtained by CPA (average PR-AUC = 0.40). The generally low performance across all pre-trained representations suggests that viability-related signals are not the dominant sources of variation in gene expression perturbations. Most representations, including PCA, performed better than direct prediction from gene expression profiles, with the exception of Biolord, which underperformed for reasons that remain unclear. On the Sci-Plex dataset, which exhibited strong generalization for gene expression reconstruction, the average PR-AUC was higher (∼0.75) compared to LINCS (average PR-AUC=0.45), suggesting that viability is more directly reflected in the transcriptomic responses (see Fig. 3b). However, comparisons with the null model revealed unexpected behavior: latent representations either matched or fell below the performance of using raw gene expression. This indicates that, while effective for reconstruction, the latent features may not capture phenotype-relevant signals in the absence of explicit supervision.

To reduce the confounding effect of basal expression level, we also evaluated models using differential expression (DE) profiles, where basal expression is subtracted to isolate drug-induced changes. In this setting, DORA performed best, achieving a PR-AUC of 0.74 and significantly outperforming Biolord, chemCPA, and raw DE inputs (*p <* 0.05), see (see Fig. SI4). This suggests that preprocessing can enhance phenotype prediction by emphasizing perturbation-specific signals. To evaluate this further, we fine-tuned the DORA model using the same 70% of labeled samples in a supervised setting, as described in Section 7. Fine-tuning led to substantial gains in phenotype prediction: the PR-AUC on the LINCS dataset improved from 0.77 to 0.86 (*p <* 0.05), and on the Sci-Plex dataset from 0.70 to 0.83 (*p <* 0.05). Fine-tuning on DE profiles also further improved performance, with DORA reaching a PR-AUC of 0.97 (see Fig. SI4). These results confirm that supervised training is essential for aligning latent features with phenotype-relevant variation.

The ability of a latent representation to support phenotype prediction is linked to how it organizes gene expression profiles in latent space. Ideally, profiles associated with similar phenotypes should map onto the same region or cluster. To investigate this, we applied an additional dimensionality reduction step using UMAP [29] to the representations benchmarked above. On the LINCS dataset, the latent representations generally reveal clustering by cell type, whereas direct gene expression inputs did not show any clear structure (see Fig. SI6). In the Sci-Plex dataset, this pattern was even more pronounced, with well-separated clusters corresponding to distinct cell types. This effect was observed across all latent models except Biolord and chemCPA, which failed to capture cell type structure (see Fig. 3c, even though Biolord already converges within the given epochs, see Fig. SI5).

With respect to viability, we found that the latent space learned by DORA on the LINCS dataset did not exhibit any discernible structure, even after fine-tuning (see Fig. SI6). This may reflect weaker alignment between viability and transcriptional response in LINCS, possibly due to noisier labels. In contrast, in the Sci-Plex dataset, fine-tuning enhanced the separation of viable and non-viable samples, suggesting that viability follows a general trend shared across all three cell types. This observation was consistent across cell types, indicating that the fine-tuned representation captured a phenotype-relevant signal that generalizes beyond individual cellular contexts.

We further examined how different models encode dose-dependent trajectories in latent space. We randomly selected 10 drug–cell pairs and visualized their trajectories across increasing dose levels. Each trajectory corresponds to a single drug–cell combination. In the Sci-Plex dataset, DORA organized these trajectories in a star-like configuration (Fig. 3c), because the control condition is anchored at the origin of the latent space. In contrast, competing methods did not exhibit a coherent structure for dose progression. A similar pattern was observed in the LINCS dataset (Fig. SI6).

These results highlight DORA’s ability to impose a consistent and interpretable organization of doseresponse dynamics in low-dimensional space. It is worth noticing that even when unsupervised latent representations can reconstruct gene expression, they do not necessarily capture phenotype-relevant signals. Supervised fine-tuning and perturbation-specific preprocessing improve alignment with viability outcomes. This indicates that the refined latent space better reflects biologically meaningful variation.

## 5 Biomarker inference using constrained architecture

Identifying key genes—biomarkers—that are associated with cancer cell growth and viability is essential for the development of targeted therapies. However, due to the complex interplay of genes within regulatory networks, inferring such biomarkers remains a significant challenge. In this section, we use the fine-tuned representation learned by DORA and its model architecture to infer candidate gene expression biomarkers linked to viability.

In our setting, a biomarker is defined as a gene whose expression contributes to predicting the viability phenotype. We therefore designed a workflow to extract, validate, and functionally characterize such genes (Fig. 4a). The approach exploits two components of the DORA architecture: (i) the matrix *D*, used to decode gene expression profiles from the latent representation *z*, and (ii) the weights *w* of the linear logistic regression classifier 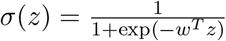. The matrix *D* represents a collection of archetypal gene expression profiles, which are linearly combined by the model to reconstruct expression profiles, similarly to PCA. To estimate how viability is influenced by gene expression, we computed the gradient of the classifier with respect to the latent representation, ∇_*z*_σ, and projected it into gene space via *VS* = *D*∇_*z*_σ. The resulting matrix *VS* assigns a weight to each gene, representing its contribution to the viability prediction, which we refer to as the viability saliency (*VS*) score. DORA tends to produce different *VS* scores across random initialization, suggesting that several biomarkers may contribute equally to the phenotype. To account for this variability, we repeat the biomarker inference 10 times and average the resulting *VS* matrices to obtain a global viability saliency score. Our inferred biomarkers are the top-*k* ranked genes according to this score.

**Figure 4:**
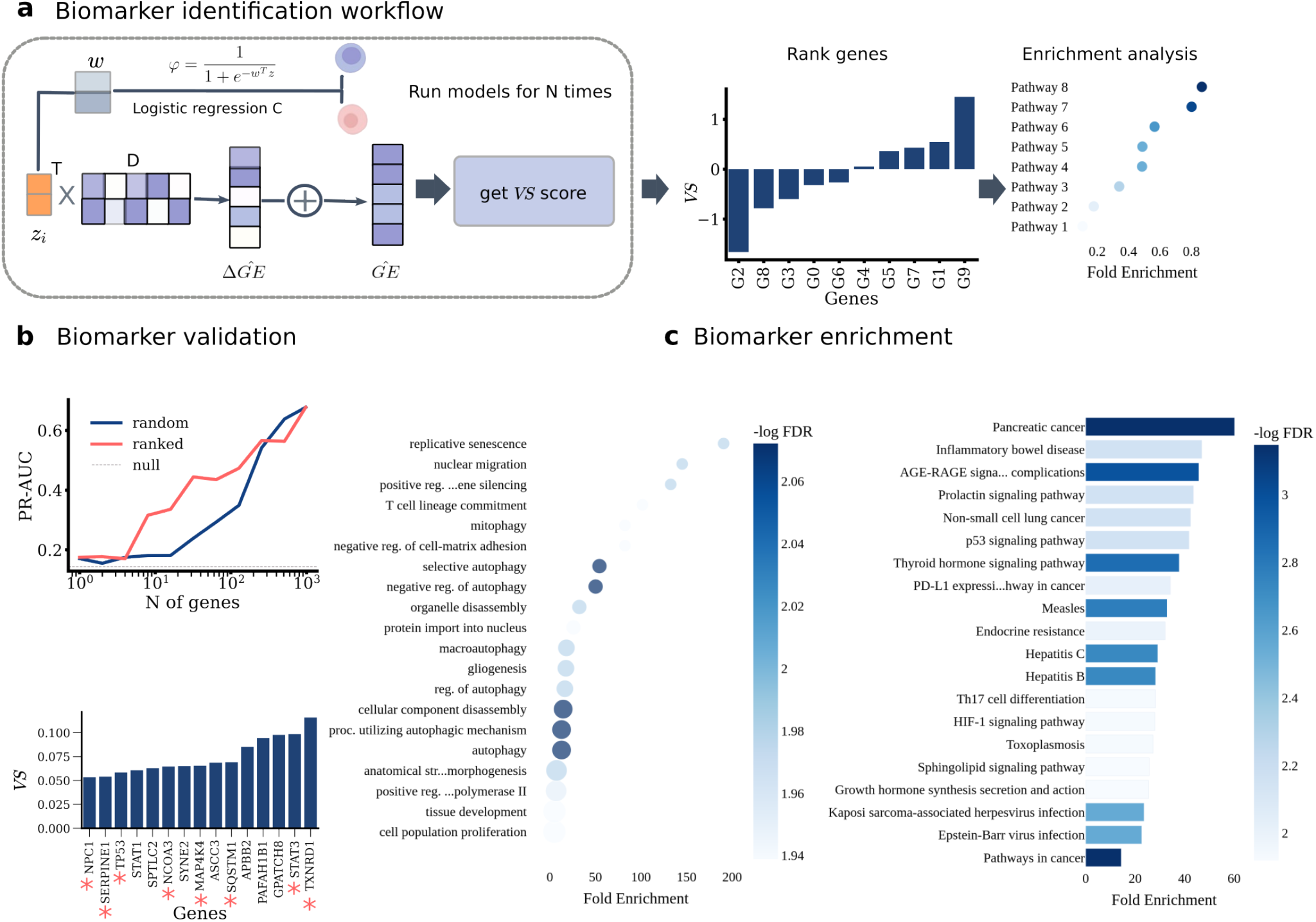
Biomarker identification using DORA and functional enrichment analysis. **a**, Workflow: DORA was trained on all gene expression (GE) data, then fine-tuned using 70% of samples with known viability. After 10 runs, average importance scores (*VS* = *D*∇_*z*_*σ*) were computed to re-rank genes; the top 20 genes were used for Gene Ontology (GO) biological process and Kyoto Encyclopedia of Genes and Genomes (KEGG) pathway enrichment. **b**, Biomarker validation: (1) PR-AUC for phenotype prediction as a function of the number of fixed genes; (2) Average *VS* scores of the top 15 genes, with blue asterisks (*) marking known cancer-related biomarkers. **c**, Enriched GO biological processes and KEGG pathways for the top 20 genes.

We next evaluated the internal consistency of this biomarker inference framework. To do so, we performed the inference on Sci-Plex to obtain average *VS* scores over 10 random initializations for the ∼900 genes retained in the study, which we ranked. The idea of our test is to compare how much of the phenotype variation is captured by DORA using either only the top-*k* ranked genes or *k* randomly selected genes. For both cases, we perturbed the non-selected genes by sampling their expression values independently from a Gaussian distribution with gene-specific means and fixed variance, removing all covariation to simulate uninformative noise. We projected the perturbed gene expression profiles into the latent space using the pseudo-inverse of *D*. Finally, we measured the viability prediction quality using a pre-trained and fine-tuned DORA instance, quantified with PR-AUC. Our results indicate that the gene ranking effectively identifies markers predictive of cell viability. The top 15 genes produced a PR-AUC of 0.35, compared to a value of 0.20 for randomly selected 15 genes. Furthermore, a consistent performance disparity existed between top-k and random-k gene subsets, with the largest difference (0.20) occurring at a gene number of around 50 (see Fig. 4b), confirming the reliability of our biomarker analysis framework.

As biomarkers, from the 978 genes retained in the analysis, we selected the top 15 ranked by the aggregated *VS* score. Notably, 8 of these were previously reported as cancer-associated, including the well-known markers TP53. TP53 plays a central role in maintaining genomic integrity and is among the most frequently mutated genes across human cancers, where it regulates cell cycle arrest, apoptosis, senescence, and DNA repair [30, 31]. The ranking was based on the averaged saliency scores across 10 random initializations, as described above, which improves stability and highlights robust associations. The recovery of established cancer genes among the highest-ranked candidates supports the biological relevance of the inferred saliency signal.

To investigate their biological roles, we performed Gene Ontology (GO) and KEGG pathway enrichment using the ShinyGO webserver [32]. We selected pathways using a false discovery rate threshold of 0.05. This threshold ensures that only statistically significant pathways are retained while accounting for the large number of gene sets tested. Then, according to fold enrichment, we chose the top 20 items. The top 5 GO terms highlighted (purple color) are selective autophagy, negative regulation of autophagy, cellular component disassembly, process utilizing autophagic mechanism and autophagy. While some of these categories are broad, they encompass key processes known to influence cancer progression and treatment response. Subcategories are provided in Table SI1 and Table SI2. The KEGG pathway analysis emphasized the pathways related to cancer. Several of the pathways were demonstrated to be highly correlated to the cancer cell growth, such as the p53 signaling pathway. Notably, p53 signaling acts as a tumor suppressor by regulating cell cycle, apoptosis, and genomic stability [33, 34], a well-known target for developing novel anti-cancer drugs. The enrichment of this pathway provides additional support that the inferred biomarkers are functionally linked to viability control in cancer cells.

To assess reproducibility, we repeated the biomarker inference procedure on the independent LINCS dataset. 11 of the 15 top-ranked genes were again documented as cancer biomarkers (see Fig.SI7). Notably, the MAPK signaling pathway was recovered in the enrichment analysis, consistent with its known role in regulating cell differentiation, proliferation, survival, and death [35]. And it has been considered as a promising therapeutic target in cancer [36]. The functional analysis details, including exact gene-pathway assignments and full ranking tables, are provided in Table SI3 and Table SI4.

These results show that we could extract interpretable biomarkers by linking viability to gene expression through the latent features generated by DORA, consistent with the recovery of known cancer genes and relevant pathways. The robustness of the predictions suggests that the method could help uncover additional biomarkers across other drugs, dosages, or cancer cell types.

## 6 Discussion

Constraining the ML architecture with biological priors can improve the prediction of transcriptomic and phenotypic signals and reduce the data required for training. The model can leverage these pre-encoded constraints to improve predictions rather than learning them. A first prior is the continuity of drug dose responses: most drug–cell pairs show no transcriptional change at low concentration, increasing progres-sively only once an effective dosage is reached. DORA explicitly encodes this by initializing from a zero latent representation and learning dose-dependent representations nonlinearly as dosage increases, unlike other methods that model concentration as a scaling factor [11, 13]. A second prior is that the effect of a drug in the latent space depends on the current latent cell state. This contrasts with additive latent models, where the drug effect remains independent of the cell state, even if the cell is already perturbed. In DORA, increasing drug dosage modulates both the magnitude and direction of the drug effect. This approach achieved state-of-the-art performance in predicting transcriptomic perturbation while relying on far fewer parameters (∼ 2 × 10^5^, see Table 3), offering a compact yet robust foundation for downstream phenotype prediction.

In both the LINCS and Sci-Plex datasets, transcriptomic profiles clustered clearly by cell type but showed no apparent separation along the viability axis. Consistent with this observation, directly predicting viability from gene expression (GE) alone yielded poor performance. It is not given either that a learned latent representation can provide a coherent representation of transcriptomes and viability at the same time. To enable viability prediction, we refined DORA in a supervised fashion by appending a linear logisticregression classifier to its architecture. This allowed us to increase the accuracy from PR-AUC of 0.77 to 0.86, well above other non-supervised models. The resulting supervised DORA analysis for Sci-Plex displayed two visible clusters of viable and non-viable samples in each cluster for cell lines. Analyses of LINCS did not display such a clearcut separation, even though viability prediction remained highly accurate. Possible explanations for differences between phenotypic separation may be due to LINCS and Sci-Plex being bulk versus single cell RNA-seq, respectively, or from the greater diversity of cell lines and drug treatments in LINCS. Finally, we note that the observed patterns may also be influenced by the choice of dimensionality reduction method. Recent benchmarking studies [37] suggest that many dimensionality reduction approaches struggle to faithfully preserve dose-dependent transcriptomic structure, which may further limit the visibility of phenotype-related organization.

As DORA successfully predicted viability from transcriptomes, this offered an opportunity to extract biomarkers for viability. This biomarker inference is enabled by the DORA decoder, which maps latent representations to their corresponding gene associations. Functionally, the decoder acts as a gene loading matrix, capturing patterns of covarying gene expression that often reflect underlying biological programs—such as those involved in cell growth regulation. Each coefficient in the decoder matrix quantifies the contribution of a gene to a particular pattern. By examining these coefficients, we were able to recover known cancer-related genes as well as propose new candidates for experimental validation. For instance, we identified TP53, a well-established tumor suppressor and key biomarker across many cancers. In addition, we discovered several less well-characterized genes that appear to participate in autophagy-related processes. This demonstrates that DORA enables interpretable predictions, relevant to actionable biological mechanisms in a clinical context.

Introducing other modelling constraints may further enhance generalization or interpretability. For example, molecular topology has been used by others to encode drug structure via graph neural networks [14], enabling the representation of novel compounds. However, such chemical priors have not yet been shown to consistently improve transcriptomic response prediction [14]. Another direction is to introduce pathway priors, since expression among genes is commonly correlated due to coordinated transcriptional regulation. This approach groups correlated genes into independent pathway blocks; therefore, the latent space could be interpreted in terms of pathways. Precedents such as PLIER [38] and DCell [39] demonstrated that pathway-aware models can capture some biological functions underlying phenotypes, such as cell growth. Therefore, extending DORA with a pathway-based latent space would be a promising next step.

Another potential extension is to apply DORA to genetic perturbation, which has become increasingly available through CRISPR-based technologies. For example, the Norman dataset contains transcriptional responses for 287 single- and double-gene perturbations [40], providing a rich resource to study the genetic interactions. Such data can serve as a proxy for identifying important genes for regulating cellular behaviors. The flexible architecture of DORA makes this extension feasible. Specifically, the drug embedding module can be replaced with gene representations derived from protein language models (e.g., ESM protein language model) [41, 42]. As DORA expands to new modalities, the next challenge is to accurately predicting unseen perturbations, which will require robust zero- and few-shot learning strategies.

## 7 Methods

### 7.1 DORA model architecture

DORA (Dose–Response Autoencoder) is a deep autoencoder framework designed to model dose-dependent transcriptomic changes and predict cell viability from latent trajectories anchored to unperturbed cellular states. The model comprises three modular components: a forward latent module, a linear decoder, and a phenotype predictor. We used ReLU activation functions all over unless specified.

Latent trajectory modeling — The model takes as input triplets (*C, D, d*), where *C* denotes a cell line, *D* a drug, and *d* a discrete dosage level. Each cell line is embedded as a fixed vector *z*_*C*_ derived from its baseline (untreated) transcriptomic profile. Each drug is embedded using ChemBERT2a [10], a pretrained transformer model on SMILES strings, to yield *z*_*D*_. The latent trajectory is initialized at *z*_0_ = (0, 0, · · ·, 0) (the unperturbed state). For each dose step, the latent state is updated via a residual MLP:

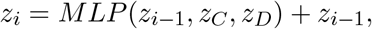

where *i* indexes increasing dose levels. The number of iterations matches the number of discrete doses observed for each (cell, drug) pair, implicitly modeling concentration escalation.

Transcriptome decoder — To preserve interpretability, the decoder was chosen as linear transforma-tion:

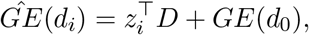

where *GE*(*d*_0_) is the unperturbed gene expression profile, *D* is a learnable weight matrix, and 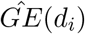 is the predicted transcriptome at dosage *d*_*i*_.

Viability fine-tuning module — Cell viability is predicted from the latent state *z*_*i*_ using a linear logistic regression classifier:

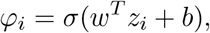

where σ denotes the sigmoid function, and *w, b* are learnable parameters.

### 7.2 Optimization of DORA

To identify the optimal chemical representations, we benchmarked 5 different molecular features: One-Hot-Encoding (OneHot), Morgan fingerprints [43], MinHashed Atom-Pair fingerprints up to a diameter of four bonds (MAP4) [44], Molecular Access System (MACCS) [45], and pre-trained representations from ChemBERTa2 (ChemBERT) [10]. We examined the generalization capacity of these representations to 9 unseen drugs. Our results showed that the choice of chemical representation had limited impact on overall prediction performance; however, ChemBERT yielded the highest reconstruction accuracy (*R*^2^ = 0.81), which was statistically superior to OneHot (*p <* 1 × 10^−5^) and MAP4 (*p <* 1 × 10^−3^). Its performance was comparable to that of MACCS (*p >* 0.05) and and Morgan fingerprints (*p >* 0.05), as shown in Fig. SI8a.

To determine the optimal dimensionality for latent representations, we conducted experiments across a range of dimensions. For Sci-Plex, we evaluated *dim* = 2, 4, 6, 8, 16, 32, 64, 128, 256, 512; for LINCS, we extended this range to include *dim* = 2, 4, 6, 8, 16, 32, 64, 128, 256, 512, 576, 640, 704, 768, 832, 896. On Sci-Plex, model performance converged at *dim* = 32, while for LINCS, convergence was observed starting at *dim* = 128, see Fig. SI8b.

The final hyperparameters for DORA are shown in Table 1 and Table 2.

**Table 1:** The hyperparameters for training DORA.

|  | LINCS | Sci-Plex |
| --- | --- | --- |
| Maximum epochs | 400 | 700 |
| Learning rate | $10^{-4}$ | $10^{-3}$ |
| Weight decay | $4 \times 10^{-5}$ | $4 \times 10^{-5}$ |
| MLP for drug dimension reduction | [384, 128] | [384, 32] |
| MLP for cell dimension reduction | [978, 128] | [977, 128] |
| Propagator | [384, 256, 128] | [192, 64, 32] |
| Decoder | [128, 978] | [32, 977] |
| Batch size | 128 | 128 |
| Early stop patience | 5 | 100 |

**Table 2:** The hyperparameters for fine-tuning DORA.

|  | LINCS | Sci-Plex |
| --- | --- | --- |
| Linear classifier $C$ | [128,1] | [32,1] |
| Learning rate for optimizer of classifier | $10^{-2}$ | $10^{-3}$ |
| Weight decay for optimizer of classifier | $10^{-6}$ | $10^{-6}$ |
| Early stop patience | 5 | 5 |
| Learning rate for optimizer of DORA | $10^{-4}$ | $10^{-5}$ |
| Weight decay for optimizer of DORA | $10^{-6}$ | $10^{-6}$ |

**Table 3:** The number of hyperparameters for models.

| Model name | N hyperparameters |
| --- | --- |
| DORA | 190,104 |
| Biolord | 152,136,107 |
| chemCPA | 1,361,687 |
| CPA | 527,574 |

### 7.3 Training of DORA

To train the model, we constructed dosage-dependent trajectories for each (cell, drug) pair by ordering GE measurements by increasing dosage. In Sci-Plex, we obtained 564 training points (188 pairs with 3 cell lines), each training points is composed with 5 points corresponding to different dosages (0, 0.01, 0.1, 1.0 and 10 *µm*). In contrast, the LINCS dataset featured greater dosage diversity, with 28 unique dosage levels and 5,064 distinct (cell, drug) combinations in the processed data.

In the first phase, the DORA model is trained on gene expression data, where each sample *GEc*_*i*_ denotes the average gene expression profile of a cell line subjected to a specific drug and dosage. The training objective minimizes the mean squared error between the observed and predicted gene expression:

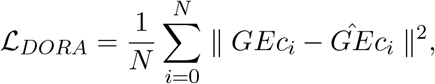

where *N* is the number of samples, 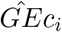 is the predicted gene expression of sample *i*. The hyperparameters that we used for training DORA are shown in Table 1.

In the second phase, DORA is fine-tuned jointly with classifier *C* using only samples annotated with phenotype labels. The classifier receives the latent representation *z*_*i*_ and outputs the predicted probability *φ*_*i*_ of inhibition. The overall objective combines the reconstruction loss with the binary cross-entropy loss:

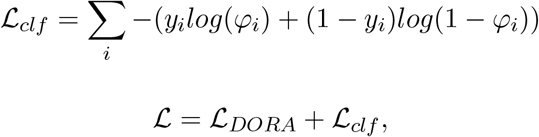

with *y*_*i*_ ∈ {0, 1}, where *y*_*i*_ = 0 denotes cell alive state and *y*_*i*_ = 1 denotes inhibited state. The hyperparameters are listed in Table 2.

In the third phase, we use the latent representations from all autoencoder-based models to predict cell viability. To ensure a fair comparison, we used a three-layer MLP for all inputs. The input layer has dimensionality equal to the input representation *dim*; the hidden layer contains 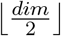 neurons; the output layer consists of 1 single neuron with a Sigmoid activation function to model the probability of cell inhibition. The models are trained using cross-entropy loss, and 10-fold cross-validation is conducted to obtain the final evaluation results.

### 7.4 Cross-dataset reference

Data preprocessing on LINCS data — To minimize experimental bias in gene expression (GE) data, we used only Phase II data (GSE70138 from GEO), specifically Level 3 profiles with 978 landmark genes from the LINCS-L1000 project [46]. The LINCS dataset represents bulk RNA expression measurements. Focusing on chemotherapy-relevant data, we retained only control and small molecule–perturbed GE profiles from Level 3. For differential expression (DE) analysis, we employed the Limma R package [47] on Level 2 data. Limma fits a linear model to account for batch effects arising from experimental time, plate wells, and other technical factors. After processing, we obtained 117,247 GE profiles and 28,801 DE profiles, covering 28 cell lines and 320 distinct drug perturbations.

Data preprocessing on Sci-Plex data — We only keep the overlapped genes (977 genes) between Sci-Plex and LINCS datasets. Later on ,we calculated the pseudo bulk gene expression for each experimental condition. After this processing, we obtained 2820 GE profiles spanning 3 cell lines, 188 drugs and 4 distinct dosages.

Mono-therapy data — DrugComb [20] has compiled a comprehensive dataset of 739,964 drug combination experiments, encompassing 8,397 unique drugs and 2,320 cell lines. For each (drugA, drugB, cell) triplet, they also measured the drug response curve for drugA or drugB treated on this specific cell line with various dosages. This resource provides valuable information on the effects of both single-drug treatments and drug combinations.

Label LINCS and Sci-Plex with viability — To label the data in LINCS and Sci-Plex, we used the monotherapy data from DrugComb [20]. We cross-referenced LINCS and DrugComb based on cell lines and drug names. We derive the inhibition rates by fitting a four-parameter dose-response model:

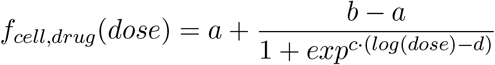

This allows us to figure out the inhibition rate for any dosages. The perturbation is labeled alive if the mapped inhibition is lower than 50% and inhibited otherwise. Overlapping (cell, drug) pairs across transcriptomic and viability datasets were retained. Final datasets:

- LINCS integrated: 4,895 samples (12.2% inhibited)
- Sci-Plex integrated: 648 samples (14.4% inhibited)

### 7.5 Biological enrichment analysis

Enrichment analysis plays a crucial role in biological data interpretation, enabling researchers to detect the overrepresented pathways for functions within a gene set. There are two widely used enrichment methods:

- Gene Ontology (GO) enrichment: It focuses on annotating genes on three categories: Biological process (BP), molecular function (MF) and cellular component (CC).
- KEGG (Kyoto Encyclopedia of Genes and Genomes) enrichment: it groups the genes to specific metabolic or signaling pathways, revealing how genes interact within biological systems.

After obtaining gene rankings using our proposed workflow, we applied ShinyGO [32] to perform GO enrichment (focusing on BP terms) and KEGG pathway analysis. We uploaded the top 20 re-ranked genes from our algorithm as input.

For GO enrichment, we reported the top 20 BP terms ranked by fold enrichment. The fold enrichment is defined as the ratio of the proportion of selected genes associated with a term to the proportion in the background gene set. The results were visualized using a bubble chart, where bubble color indicates the false discovery rate (FDR)—with deeper purple representing lower p-values—and bubble size corresponds to the number of genes associated with each term. The Y-axis lists the enriched BP terms, and the X-axis shows the fold enrichment score.

For KEGG enrichment, we reported the top 10 pathways using a bar plot. Bar color indicates the FDR, the Y-axis shows the pathway names, and the X-axis displays fold enrichment scores.

In both analyses, we placed particular emphasis on items with deep purple coloring, which correspond to lower FDR values and therefore higher statistical significance—indicating that the enrichment is less likely to have occurred by chance.

### 7.6 Computational resources

All models were trained on NVIDIA RTX A5000. DORA has ∼ 1 × 10^5^ learnable parameters. Training time per dataset was about 5 minutes. The number of hyperparameters for each model is listed here:

### 7.7 Evaluation protocols

Prediction of drug-induced transcriptomics — Performance was measured using the coefficient of determination *R*^2^ between predicted and observed gene expression. This quantifies how well model predictions recapitulate measured gene expression values. *R*^2^ was computed both across all genes and for the top 50 most highly differentially expressed genes (DEGs). Three evaluation settings were used:

- Random split: Samples were randomly partitioned into 80% training, 10% validation, and 10% test sets. This split evaluates in-distribution performance under standard conditions, where the model observes overlapping cell lines and drugs during training and testing.
- Unseen compounds extrapolation: A challenging holdout setup designed to test generalization to previously unseen drugs. We held out entire drugs during training (including all their doses and cellline combinations) and evaluated only on these unseen compounds. This setup mimics real-world drug discovery, where models must predict responses to novel molecules never encountered during training. Following the previous studies [11, 14, 15], we held out 9 drugs (Dacinostat, Givinostat, Belinostat, Hesperadin, Quisinostat, Alvespimycin, Tanespimycin, TAK-901, and Flavopiridol) that covers different targeted pathways.
- Unseen cell extrapolation: A cross-context generalization setup that tests the ability to predict responses in unseen cell types. For each cell line in the dataset, we held out all samples of that cell line during training (including all drugs and doses) and evaluated performance exclusively on the held-out cell type. This assesses whether the model learns generalizable perturbation rules rather than cell-line-specific patterns.

We compared 3 state-of-the-art methods and two baselines:

- CPA [11]: A disentangled autoencoder that decomposes perturbed gene expression into independently optimized cell and drug embeddings using adversarial training. It models transcriptional responses as an addition of cell, drug, and perturbation attributes. The drug and cell representations should be learned from data itself.
- chemCPA [14]: An extension of CPA that replaces one-hot drug encoding with graph neural network (GNN)-based molecular embeddings to improve generalization to unseen compounds, while retaining the additive latent framework of CPA. This allows autoencoder to predict on new compounds.
- Biolord [15]: A disentangled representation model that learns separate biological attributes (e.g., molecular, genetic) and concatenates them nonlinearly in the decoder. The representations for attributes are learned in a supervised manner.
- Mean expression: A non-parametric reference model that compute the average expression value of each gene across all samples in the training set. This model captures global expression trends, serving as a measure of how much predictive signal is captured beyond simple population averages.
- Identity (untreated) model: A minimal control model that always returns the unperturbed (control) gene expression profile for any input condition, regardless of cell type, drug, or dosage. It assumes no transcriptional change occurs upon drug treatment and establishes a lower performance bound for evaluating whether a model captures biologically meaningful perturbation responses.
- CPA, ChemCPA, and Biolord were implemented using official open-source code with default hyperparameters for fair comparison.

Viability prediction — Latent representations from each model were used to train a shared 3-layer MLP classifier. Performance was reported as precision–recall AUC (PR-AUC). This assesses the model’s capacity in the prediction of inhibited cell states.

Statistical evaluation — Statistical comparisons between models were performed using t-tests to assess whether one model yielded a significantly higher mean *R*^2^ than another.

## 8 Data availability

Datasets in this paper are publicly available. The LINCS dataset can be downloaded from: https://www.ncbi.nlm.nih.gov/geo/query/acc.cgi?acc=GSE70138. The original datasets of Sci-Plex can be downloaded from: https://f003.backblazeb2.com/file/chemCPA-datasets/sciplex_complete_middle_subset.h5ad. The data from DrugComb can be obtained from: https://drugcomb.org/download/. However, to get the drug response curve, we need to use a data mining method from the API of DrugComb.

The preprocessed data can be obtained by using the notebook in our GitHub.

## 9 Code availability

All the code is on the GitHub: https://github.com/LBiophyEvo/dora_reproducibility.git. DORA is provided in the format of a Python (version 3.10) package.

