## Supplemental Information for "DORA: a dose-response autoencoder for interpretable transcriptome-to-viability prediction"

### Supplementary Information

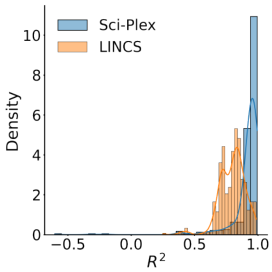


Fig. SI1: Comparison of the $R^2$ distribution between LINCS and Sci-Plex datasets. The $R^2$ values represent the similarity between control and perturbed gene expression profiles for all validation samples under the random split scenario.

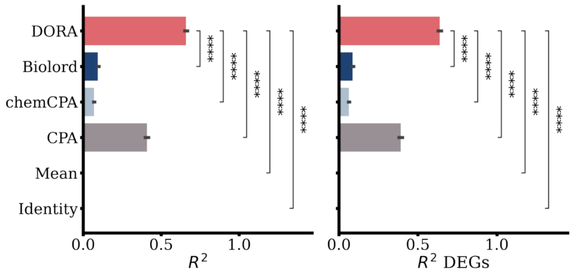


Fig. SI2: Prediction of differential gene expression using DORA and other state-of-the-art models (CPA, chemCPA, Biolord) on LINCS data. Differential gene expression is obtained by Limma. Left panel: Box plots of $R^2$ scores across all genes. Right panel: Box plots of $R^2$ scores for the top 50 differentially expressed genes (DEGs). t-test is used to assess if the performances between models are statistically significant (****: $p \leq 10^{-4}$).

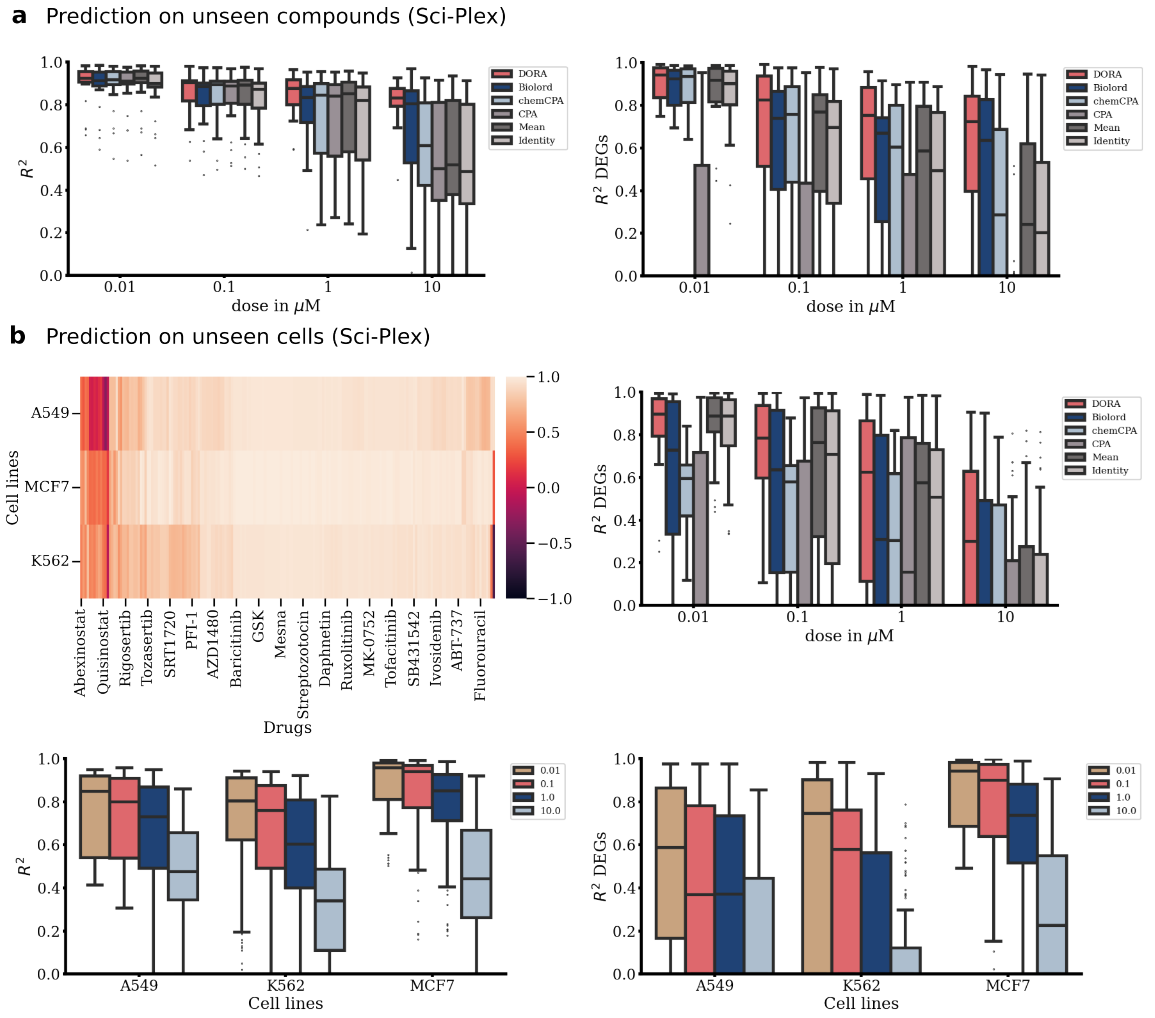


Fig. SI3: Reconstruction of gene expression (GE) profiles in the Sci-Plex dataset for unseen drugs and unseen cell lines. **a**, GE prediction performance for unseen drugs across different models. $R^2$ scores are stratified by treatment dosage. Left, box plots of $R^2$ values across all genes; right, $R^2$ scores for the top 50 differentially expressed genes (DEGs). **b**, GE prediction for unseen cell lines. First panel, $R^2$ values quantifying similarity between control and perturbed GE profiles across all (cell line, drug) combinations following 10 $\mu m$ treatment, demonstrating that most perturbations do not induce substantial GE changes across the three cell lines. Second panel, GE prediction performance of different models, with $R^2$ scores for DEGs stratified by model and dosage. Third and fourth panels, performance of the DORA model, stratified by cell line and treatment dosages.

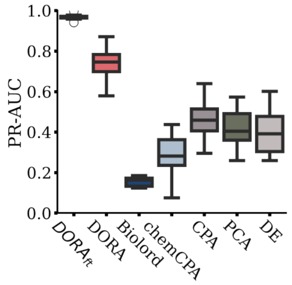


Fig. SI4: Prediction of phenotype by using differential gene expression of LINCS data. The X-axis indicated the latent representation from different models. The Y-axis shows the average PR-AUC by using 10-fold cross-validation.

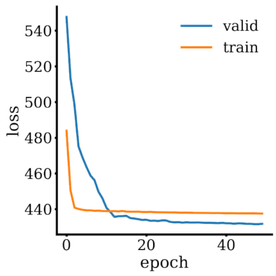


Fig. SI5: The evolution of reconstruction loss on training and validation of Biolord model.

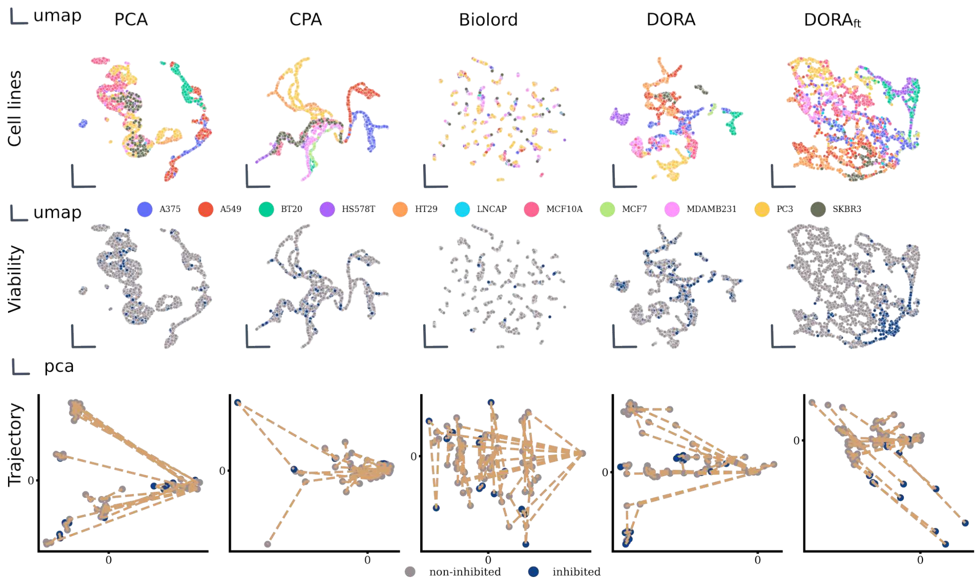


Fig. SI6: UMAP projections of LINCS colored by cell line (hue) and phenotype (lightness: dark = inhibited, light = alive). GE uses raw expression data; PCA keeps the top 100 components, and others use model-derived embeddings.

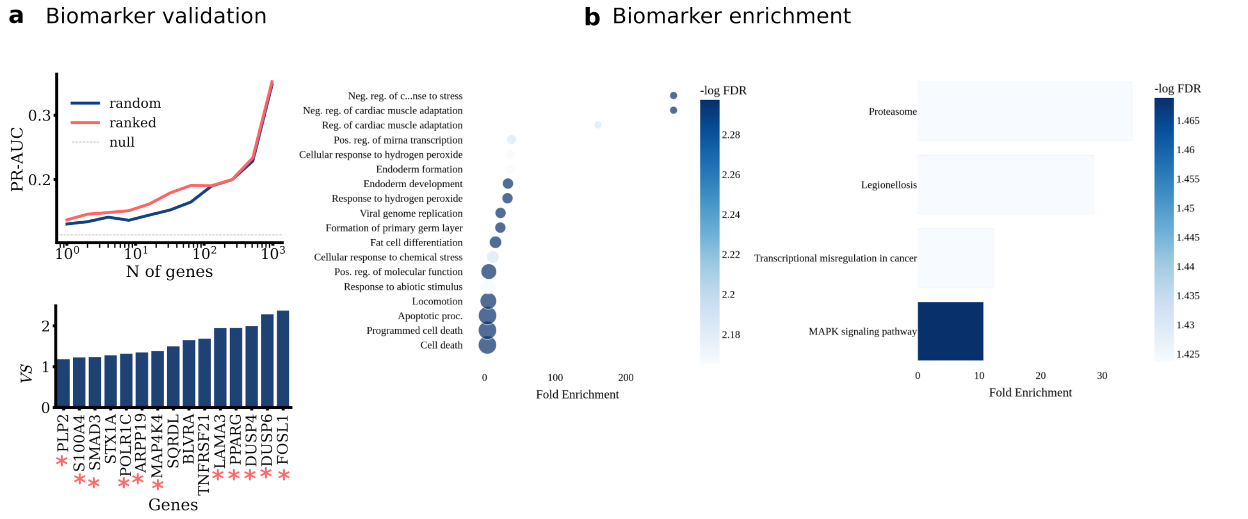


Fig. SI7: Results of biomarker identification on LINCS GE data. Left panel: PR-AUC for phenotype prediction as a function of the number of fixed top-ranked genes, demonstrating their predictive importance. Second panel: Average $VS$ scores of the top 15 genes; blue asterisks (*) mark known cancer-related biomarkers. Right two panels: Enriched GO biological processes and KEGG pathways for the top 20 genes.

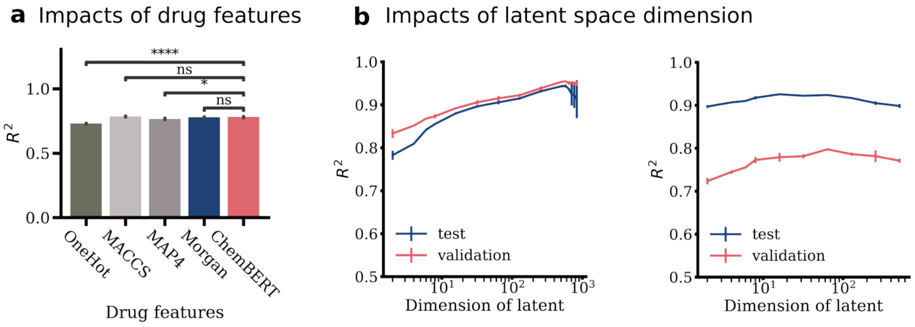


Fig. SI8: Impact of drug features and latent dimensionality on gene expression (GE) prediction. **a**, Comparison of GE prediction performance on unseen drugs in Sci-Plex using different drug feature encodings: OneHot, MACCS, MAP4, Morgan, and ChemBERT. Significance: ns $(0.05 < p \leq 1)$, * $(0.01 < p \leq 0.05)$, **** $(p \leq 0.0001)$. **b**, Effect of latent dimension size on GE prediction accuracy ($R^2$) for the LINCS and Sci-Plex datasets. X-axis, number of latent dimensions; Y-axis, $R^2$ values on validation and test sets.

Table SI1: Biological enrichment for top 20 genes in Sciplex.

| Enrichment FDR | nGenes | Fold Enrichment | Pathways | Genes |
|---|---|---|---|---|
| 0.008 | 5 | 12.9 | autophagy | SQSTM1 SPTLC2 STAT3 NPC1 TP53 |
| 0.008 | 3 | 50.3 | negative reg. of autophagy | STAT3 NPC1 TP53 |
| 0.008 | 5 | 14.7 | cellular component disassembly | SQSTM1 ASCC3 PAFAH1B1 TP53 MAP4K4 |
| 0.008 | 3 | 54.5 | selective autophagy | SQSTM1 SPTLC2 TP53 |
| 0.008 | 5 | 12.9 | proc. utilizing autophagic mechanism | SQSTM1 SPTLC2 STAT3 NPC1 TP53 |
| 0.011 | 2 | 145.3 | nuclear migration | PAFAH1B1 SYNE2 |
| 0.011 | 4 | 16.5 | reg. of autophagy | SPTLC2 STAT3 NPC1 TP53 |
| 0.011 | 4 | 18.2 | macroautophagy | SQSTM1 SPTLC2 NPC1 TP53 |
| 0.011 | 4 | 17.7 | gliogenesis | PAFAH1B1 SYNE2 TP53 STAT3 |
| 0.011 | 6 | 7.4 | anatomical structure formation involved in morphogenesis | SERPINE1 PAFAH1B1 TP53 STAT3 TXNRD1 STAT1 |
| 0.011 | 2 | 132.6 | positive reg. of post-transcriptional gene silencing | TP53 STAT3 |
| 0.011 | 2 | 190.7 | replicative senescence | SERPINE1 TP53 |
| 0.011 | 3 | 32.5 | organelle disassembly | SQSTM1 ASCC3 TP53 |
| 0.011 | 6 | 7.0 | positive reg. of transcription by RNA polymerase II | NCOA3 STAT1 STAT3 TP53 APBB2 SQSTM1 |
| 0.012 | 2 | 82.5 | mitophagy | SQSTM1 TP53 |
| 0.012 | 2 | 82.5 | negative reg. of cell-matrix adhesion | SERPINE1 MAP4K4 |
| 0.012 | 2 | 101.7 | T cell lineage commitment | TP53 STAT3 |
| 0.012 | 3 | 26.0 | protein import into nucleus | STAT3 TP53 SQSTM1 |
| 0.012 | 7 | 5.0 | cell population proliferation | STAT1 STAT3 TP53 PAFAH1B1 SQSTM1 TXNRD1 ASCC3 |
| 0.012 | 7 | 5.1 | tissue development | SERPINE1 PAFAH1B1 SPTLC2 TP53 TXNRD1 STAT1 NCOA3 |

Table SI2: KEGG enrichment for top 20 genes in Sci-Plex.

| Enrichment FDR | nGenes | Fold Enrichment | Pathway | Genes |
|---|---|---|---|---|
| 0.001 | 5 | 14.4 | Pathways in cancer | STAT1 STAT3 TP53 TXNRD1 NCOA3 |
| 0.001 | 3 | 60.2 | Pancreatic cancer | STAT1 STAT3 TP53 |
| 0.001 | 3 | 45.8 | AGE-RAGE signaling pathway in diabetic complications | SERPINE1 STAT1 STAT3 |
| 0.001 | 3 | 37.8 | Thyroid hormone signaling pathway | STAT1 TP53 NCOA3 |
| 0.002 | 3 | 32.9 | Measles | STAT1 STAT3 TP53 |
| 0.002 | 3 | 29.1 | Hepatitis C | STAT1 STAT3 TP53 |
| 0.002 | 3 | 28.2 | Hepatitis B | STAT1 STAT3 TP53 |
| 0.003 | 3 | 23.6 | Kaposi sarcoma-associated herpesvirus infection | STAT1 STAT3 TP53 |
| 0.003 | 3 | 22.7 | Epstein-Barr virus infection | STAT1 STAT3 TP53 |
| 0.007 | 2 | 41.8 | p53 signaling pathway | SERPINE1 TP53 |
| 0.007 | 2 | 43.6 | Prolactin signaling pathway | STAT1 STAT3 |
| 0.007 | 2 | 42.4 | Non-small cell lung cancer | STAT3 TP53 |
| 0.007 | 2 | 46.9 | Inflammatory bowel disease | STAT1 STAT3 |
| 0.010 | 2 | 34.3 | PD-L1 expression and PD-1 checkpoint pathway in cancer | STAT1 STAT3 |
| 0.010 | 2 | 32.1 | Endocrine resistance | TP53 NCOA3 |
| 0.012 | 2 | 28.0 | HIF-1 signaling pathway | SERPINE1 STAT3 |
| 0.012 | 2 | 28.2 | Th17 cell differentiation | STAT1 STAT3 |
| 0.012 | 2 | 27.2 | Toxoplasmosis | STAT1 STAT3 |
| 0.012 | 2 | 25.6 | Sphingolipid signaling pathway | TP53 SPTLC2 |
| 0.012 | 2 | 25.4 | Growth hormone synthesis secretion and action | STAT1 STAT3 |

Table SI3: Biological process enrichment for top 20 genes in LINCS.

| Enrichment FDR | nGenes | Fold Enrichment | Pathway | Genes |
|---|---|---|---|---|
| 0.005 | 2 | 267.3 | Neg. reg. of cardiac muscle adaptation | PPARG SMAD3 |
| 0.005 | 2 | 267.3 | Neg. reg. of cardiac muscle hypertrophy in response to stress | PPARG SMAD3 |
| 0.007 | 2 | 160.4 | Reg. of cardiac muscle adaptation | PPARG SMAD3 |
| 0.007 | 3 | 38.2 | Pos. reg. of mirna transcription | PPARG SMAD3 FOSL1 |
| 0.007 | 3 | 35.9 | Cellular response to hydrogen peroxide | NET1 HSPA8 BNIP3 |
| 0.007 | 3 | 35.4 | Endoderm formation | DUSP4 LAMA3 HMGA2 |
| 0.005 | 4 | 33.1 | Endoderm development | DUSP4 SMAD3 LAMA3 HMGA2 |
| 0.005 | 4 | 32.4 | Response to hydrogen peroxide | NET1 HSPA8 FOSL1 BNIP3 |
| 0.005 | 4 | 22.6 | Viral genome replication | HMGA2 HSPA8 PPIE PLSCR1 |
| 0.005 | 4 | 22.3 | Formation of primary germ layer | DUSP4 SMAD3 LAMA3 HMGA2 |
| 0.005 | 5 | 15.4 | Fat cell differentiation | SMAD3 PPARG BNIP3 PSMB8 HMGA2 |
| 0.007 | 5 | 11.7 | Cellular response to chemical stress | CAB39 NET1 HSPA8 PPARG BNIP3 |
| 0.005 | 8 | 6.0 | Pos. reg. of molecular function | DNAJB1 PPARG CAB39 HMGA2 PLSCR1 SMAD3 NET1 MAP4K4 |
| 0.007 | 8 | 5.3 | Response to abiotic stimulus | CAB39 DNAJB1 NET1 HSPA8 PPARG FOSL1 BNIP3 SMAD3 |
| 0.005 | 9 | 5.3 | Locomotion | HSPA8 PPARG LAMA3 SLC37A4 SMAD3 S100A4 MAP4K4 PLP2 FOSL1 |
| 0.005 | 11 | 4.2 | Apoptotic proc. | DUSP6 TNFRSF21 BNIP3 PPARG HMGA2 NET1 PLSCR1 HSPA8 SMAD3 FOSL1 MAP4K4 |
| 0.005 | 11 | 4.0 | Programmed cell death | DUSP6 TNFRSF21 BNIP3 PPARG HMGA2 NET1 PLSCR1 HSPA8 SMAD3 FOSL1 MAP4K4 |
| 0.005 | 11 | 4.0 | Cell death | DUSP6 TNFRSF21 BNIP3 PPARG HMGA2 NET1 PLSCR1 HSPA8 SMAD3 FOSL1 MAP4K4 |

Table SI4: KEGG enrichment for top 20 genes in LINCS.

| Enrichment FDR | nGenes | Fold Enrichment | Pathway | Genes |
|---|---|---|---|---|
| 0.038 | 2 | 34.9 | Proteasome | PSMB8 PSME2 |
| 0.038 | 2 | 28.6 | Legionellosis | HSPA8 BNIP3 |
| 0.038 | 3 | 12.3 | Transcriptional misregulation in cancer | DUSP6 PPARG HMGA2 |
| 0.034 | 4 | 10.7 | MAPK signaling pathway | DUSP4 DUSP6 HSPA8 MAP4K4 |
